# A short intrinsically disordered domain of MCPyV ALTO regulates host TBK1 signaling and MCPyV latency

**DOI:** 10.1101/2025.08.09.669491

**Authors:** Taylor E. Senay, Xiaomei Li, Sneha G. Shirhattikar, Tiana T. Luo, Jianxin You

## Abstract

Merkel Cell Polyomavirus (MCPyV) is an oncogenic human polyomavirus that latently infects most adults. Although the causative link between MCPyV and Merkel Cell Carcinoma (MCC) is well established, the molecular mechanisms that govern viral latency and prevent oncogenic progression remain poorly understood. We previously reported that the MCPyV early region protein ALTO is a key modulator of the STING-TBK1 signaling axis, enabling the virus to co-opt host innate immune pathways to suppress excessive viral replication and promote latency over transformation. In this study, we expand on this model by identifying a short, essential domain within ALTO that is required for TBK1 activation. This domain, which we term LIT (Lost in Tau), is necessary for ALTO-TBK1 interaction but dispensable for ALTO trafficking and its interactions with STING or Src. When expressed alone, the LIT domain functions as a dominant negative inhibitor of wild-type ALTO, competitively blocking TBK1 activation through a novel TBK1 interaction domain. Deletion of the LIT domain from ALTO not only abolishes TBK1 interaction and downstream phosphorylation but also eliminates TBK1-mediated suppression of MCPyV replication during early infection of human dermal fibroblasts (HDFs). These findings provide mechanistic insight into how ALTO promotes viral persistence and immune evasion. More broadly, they highlight the functional importance of intrinsically disordered regions in modulating host–virus interactions and suggest that MCPyV latency is actively maintained through a finely tuned balance of pro- and anti-viral signaling. Targeting domains such as LIT may offer new strategies for regulating TBK1 activity or disrupting viral persistence.

**Importance:** Merkel cell polyomavirus causes lifelong, latent infections in the skin of most people. When this latency is perturbed, the virus can give rise to Merkel Cell Carcinoma, an aggressive and difficult-to-treat skin cancer. Efforts to prevent this cancer depend on understanding what controls viral latency and persistence. We previously reported that MCPyV stimulates the host’s STING-TBK1 signaling axis to limit its own replication. In this work, we identify and characterize a short amino acid motif within the ALTO protein’s intrinsically disordered region that is required for this immune-stimulating activity. This region appears to be critical for helping the virus maintain latency by fine-tuning the host’s response. Our findings provide new insight into MCPyV’s latency mechanism and may help guide future approaches to prevent or treat Merkel cell carcinoma by targeting viral or cellular factors involved in long-term infection.

## Introduction

The viral family *Polyomaviridae* includes approximately 117 species of non-enveloped viruses carrying small (∼5 kb) circular double-stranded DNA genomes(1). Of these, around 16 are thought to be human-specific(2), and only one, Merkel cell polyomavirus (MCPyV or MCV), has been conclusively demonstrated to be carcinogenic(3). MCPyV is ubiquitous in the human population and, for most individuals, exists as a silent member of the skin virome, establishing lifelong episomal latency in cutaneous cells(4–8). However, in a subset of cases, MCPyV integrates into the host genome, where it drives oncogenesis of the rare but aggressive neuroendocrine skin cancer Merkel cell carcinoma (MCC) through constitutive high-level expression of its Small Tumor (sT) antigen and a truncated form of its Large Tumor (LT) antigen(3, 9, 10).

Nearly two decades after its discovery as the major causative agent in approximately 80% of MCCs(3, 11, 12), most mechanistic studies of MCPyV have focused on LT and sT. These viral effectors have homologs in other well-characterized polyomaviruses, such as SV40, and their roles in MCPyV replication and transformation are well established(13–15). However, the upstream events that initiate viral genome truncation and integration, as well as the cellular conditions that distinguish persistent infection from oncogenic progression, remain poorly understood. These represent critical gaps in our knowledge of MCPyV biology. Elucidating the regulatory mechanisms that govern this transition is essential for improving early detection and prevention of MCC, which currently has limited and often non-durable treatment options.

There is currently no consensus on how MCPyV establishes, maintains, or exits latency. Clinical observations reveal that immunosuppressed individuals carry higher MCPyV viral burdens and are at substantially increased risk of developing MCC, suggesting that innate immune control is a key determinant of viral latency(16–18). The cyclic GMP-AMP synthase–stimulator of interferon genes (cGAS–STING) pathway, a cytosolic DNA-sensing system that activates type I interferon and proinflammatory signaling, has emerged as particularly important in this context(19–25). We have previously reported that STING expression is dramatically reduced in virus-positive MCC tumors, and that reactivation of STING signaling promotes MCC cell death and T-cell infiltration(26). Conversely, during early infection, MCPyV strongly activates the STING pathway and its downstream proinflammatory effectors, which in turn restricts viral replication and supports the establishment of a balanced, persistent infection(27–29). Together, these findings suggest that MCPyV latency is not a passive process, but rather one that is tightly regulated through interactions between viral effectors and host innate immune pathways. Disruption of this equilibrium, whether through viral mutation or host immune suppression, may be a critical inflection point that drives malignant progression.

While LT and sT have been the primary focus of MCPyV research, less attention has been paid to another early region protein, the Alternative Large Tumor antigen open reading frame (ALTO). First identified in 2014, ALTO was initially thought to be dispensable for replication in transfection-based assays(30). However, recent studies, including our own, have revealed its active role in immune signaling. ALTO has been shown to modulate Src-family kinases, PLCγ1 signaling(31), and the NFκB pathway(31, 32). In our recent work, we demonstrated that ALTO acts as a modulator of the STING–TBK1 axis, forming a signaling complex with STING, TBK1, and Src to induce TBK1 autophosphorylation, leading to its degradation and resulting in viral replication suppression and latency promotion(29). This function places ALTO in the unique position of acting as an “anti-viral viral protein” that dampens its own replication evade immune detection and persist in the host.

These findings reshape our understanding of ALTO as a multifunctional effector in MCPyV biology, one that fine-tunes host responses through structurally flexible, context-dependent interactions. In this study, we build upon our prior work to dissect the structure-function relationship of ALTO in greater detail. Using a systematic truncation approach, we identify a short, previously uncharacterized region within ALTO’s intrinsically disordered domain that is required for its interaction with TBK1 and its immunomodulatory function. Through this analysis, we aim to illuminate how ALTO contributes to latency maintenance, how disruption of its function may promote oncogenesis, and how ALTO-targeted mechanisms could be leveraged to better understand or intervene in MCPyV-associated disease.

## Results

### Identification of the TBK1-activating domain within ALTO’s N-terminal intrinsically disordered region

As shown in our previous study published in 2024, one of ALTO’s key features is its ability to stimulate TBK1 autophosphorylation(29). To identify the region(s) of ALTO responsible for stimulating TBK1 autophosphorylation, we began by evaluating the current understanding of ALTO’s structural features. The ALTO protein remains poorly characterized at the structural level, and even its precise length is debated: while early studies described a 250-amino acid (aa) product encoded across two exons, more recent annotations favor a 248-aa version encoded within a single exon(30–32). As no conclusive functional differences between these isoforms have been reported, we included both in our *in vitro* studies and observed no phenotypic distinctions between them.

Multiple reports of *in silico* analyses agree that ALTO’s most stable structural feature is a C-terminal alpha helix (approximately aa 200-250) that contains a 22-aa transmembrane or hydrophobic domain (aa 225-246). In contrast, the N-terminal region, representing roughly 80% of the protein, is intrinsically disordered (Fig. 1A)(33–35). Although intrinsically disordered regions (IDRs) are known to be highly plastic and capable of a wide range of functions(36–39) their lack of fixed globular shape makes structure-based functional prediction based on modeling difficult. We therefore took a methodical but exploratory approach using progressive N-terminal focused truncation. Each truncation shortened the protein by 30-aa compared to the one before, allowing us to examine region-specific phenotypic variation with fine resolution (Fig. 1B). In addition to the progressive N-terminal truncation mutants, we constructed a C-terminal ALTO truncation variant lacking the hydrophobic or transmembrane domain (ALTOΔTM). These variant sequences were generated using the NanoBiT platform, a structural complementation-based split enzyme system which quantifies tagged protein-protein interactions in live cells via relative luminescence. We have previously found that ALTO favors N-terminal functional tags(29), so all constructs were generated with N-terminal LgBiT and SmBiT tags—the large and small subunits, respectively, of a reconstitutable NanoLuc luciferase enzyme—enabling protein detection and downstream protein-protein interaction analysis (Fig. 1B).

**Figure 1:**
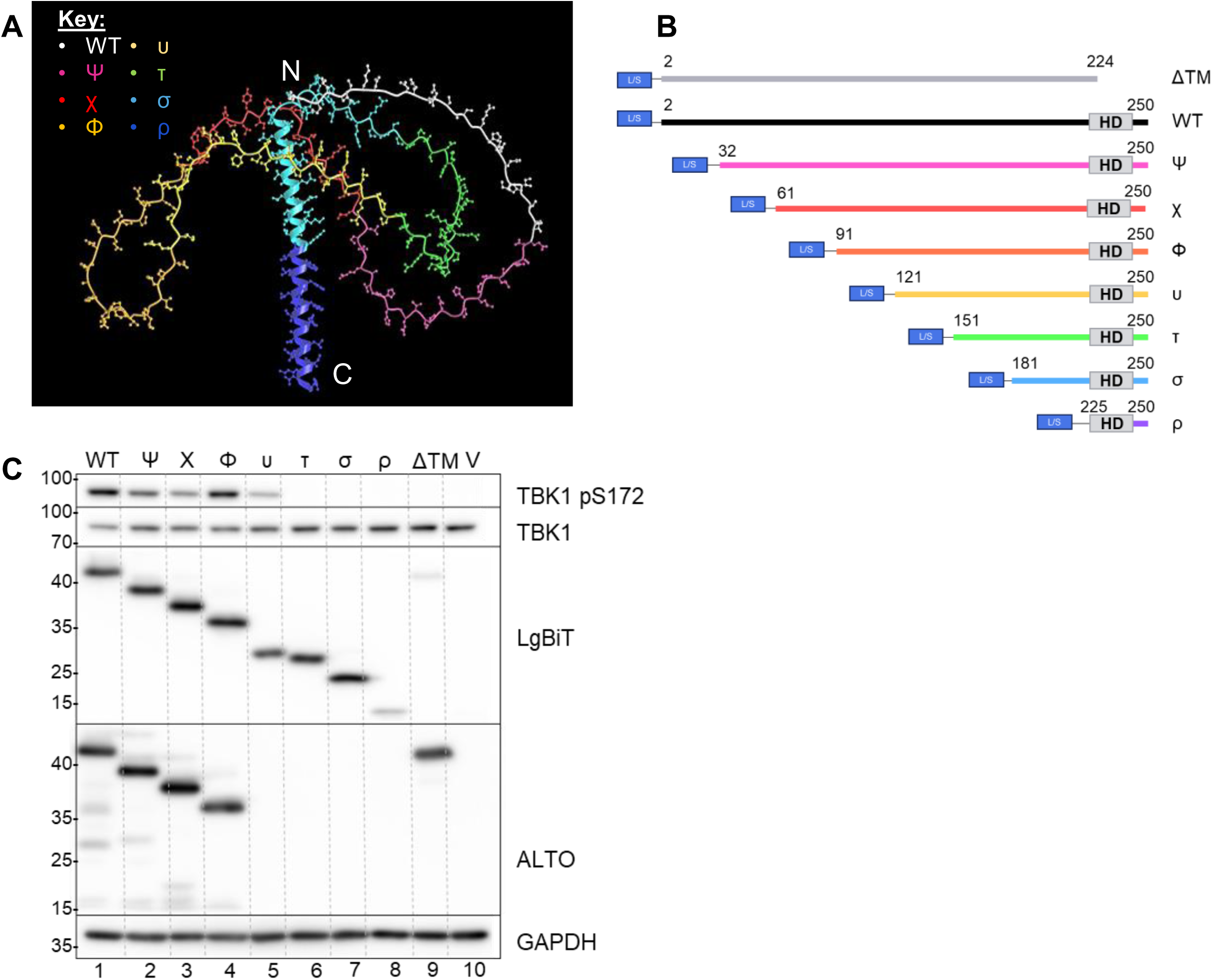
Progressive N-terminal truncation reveals ALTO functional regions. (A) Predicted tertiary structure model of ALTO (AlphaFold2 Colab) with truncation mutant lengths highlighted. Chain colors correspond to the maximum N-terminal length of their respective mutant, with each mutant consisting of its own color portion and all subsequent C-terminal portions. (B) Schematic diagram of LgBiT- and SmBiT-(L/S) tagged progressive N-terminal ALTO truncation mutants. The hydrophobic or transmembrane (TM) domain is indicated as “HD”. (C) Western blots of HEK 293T lysates collected 24 hours post-transfection with 2ug of the indicated LgBiT-tagged progressive ALTO truncation mutant (or an empty vector control, V). Primary antibodies are listed on the right.

To test which region or regions of ALTO govern its TBK1-stimulatory activity, we transfected HEK 293T cells with constructs encoding LgBiT-tagged ALTO progressive truncation mutants or an empty vector control. Expression of most variants was roughly equivalent, and all migrated to positions consistent with their predicted molecular weights (Fig. 1C). Although structural prediction of ALTO remains largely inconclusive, AlphaFold modeling suggested that all variants preserved the characteristic N-terminal disordered region and C-terminal alpha helix (Fig. S1). Since ALTO is also known to form homo-oligomers(29, 31), we used ALTO-ALTO interactions as a functional readout to assess the structural integrity of the truncation mutants. Self-interaction assays confirmed that variants retained appropriate homo- and hetero-oligomerization capability (Fig. S2), further supporting the functional integrity of these constructs.

We then tested whether the ALTO truncation mutants retained the ability to stimulate TBK1 autophosphorylation. As observed with wildtype (WT) ALTO, the first four truncation mutants (ALTOΨ, ALTOχ, ALTOΦ, and ALTOυ) retained the ability to stimulate TBK1 autophosphorylation (Fig. 1C, lanes 1-5). In contrast, the ALTOτ truncation (aa 151–250) and all shorter N-terminal truncated variants completely lost this activity despite comparable protein expression levels (Fig. 1C, lane 6-8). ALTOΔTM also failed to stimulate TBK1 autophosphorylation and was expressed at a relatively lower level (Fig. 1C, lane 9). This, taken with its overall lowered homo- and hetero-oligomerization capability (Fig. S2) supports previous reporting that the putative transmembrane or hydrophobic domain is required for robust expression of ALTO(32).

### The LIT domain is specifically required for ALTO-TBK1 interaction but not for binding STING or Src

The loss of TBK1 autophosphorylation in ALTOτ and shorter variants may indicate a disruption in the formation of the ALTO-STING-TBK1-Src “signal complex” we previously described(29). To determine whether the loss of TBK1-stimulatory activity in certain truncation mutants was due to disrupted complex formation—and to identify which specific protein-protein interactions might underlie the observed loss of function—we co-transfected HEK 293T cells with SmBiT-tagged ALTO truncation mutants and LgBiT-tagged STING, TBK1, or Src (Fig. 2A-C). Kinetic monitoring of these interactions showed that after a short equilibration period, all readings remained relatively stable over the observation window; therefore, single representative timepoints are presented throughout this study (Fig. S3). For all three interactors, interaction strength was substantially reduced or eliminated at different truncation points, suggesting that STING, Src, and TBK1 each rely on distinct ALTO regions for binding.

**Figure 2:**
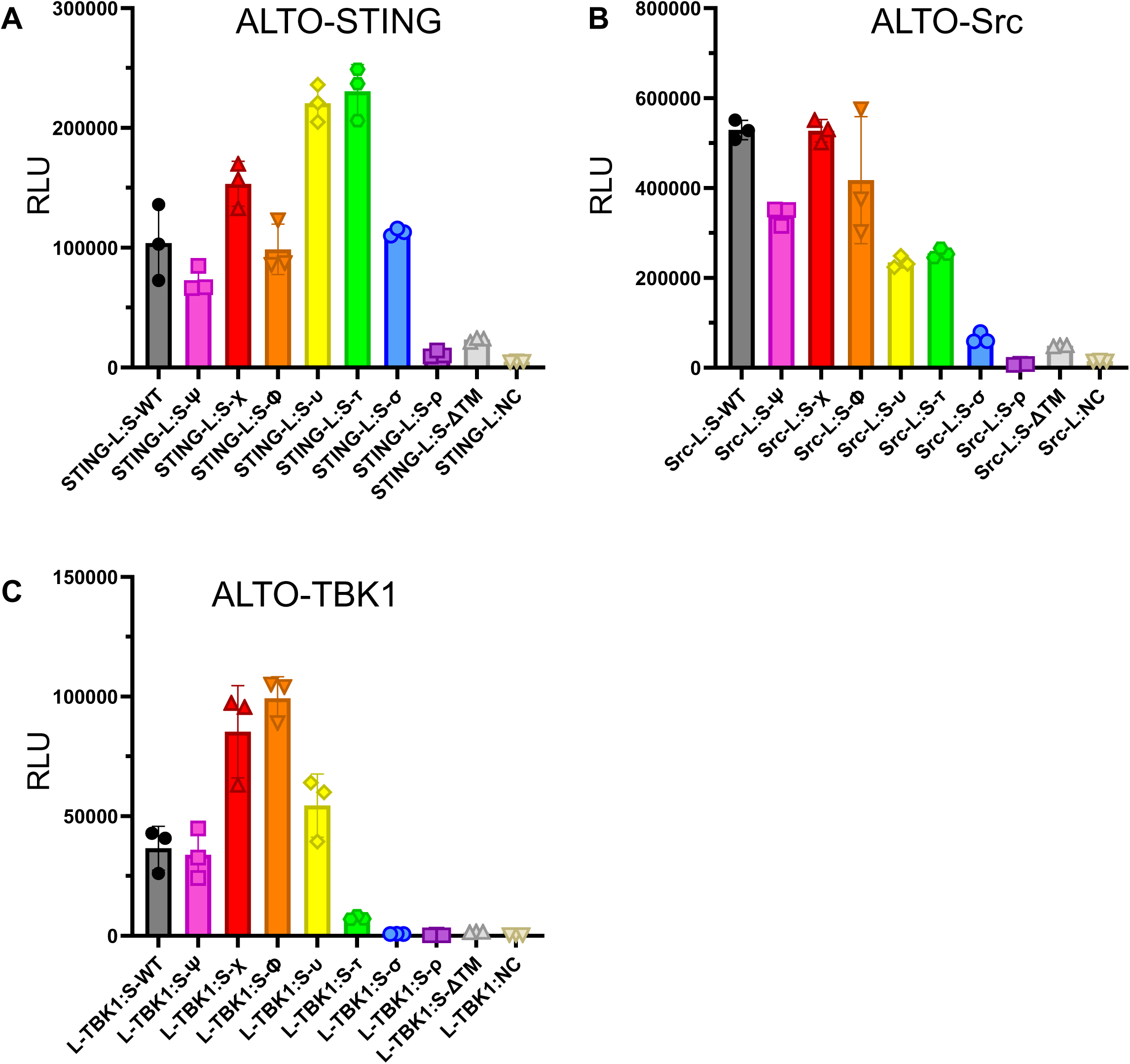
ALTOτ is markedly deficient in TBK1 interaction. (A-C) Luminescence readings of HEK 293T cells transfected with pairs of plasmids carrying NanoBiT-tagged ALTO progressive truncation mutants and either (A) STING, (B) Src, and (C) TBK1, or the Small BiT negative control construct (NC). Readings were taken at 20 hours (ALTO-TBK1) or 24 hours (ALTO-STING and ALTO-Src) post-transfection. Points indicate biological replicate wells, bars indicate means, and error bars indicate standard deviations. L-protein and protein-S (and similar) indicate the named protein tagged with the LgBiT at its N-terminus or SmBiT at its C-terminus, respectively.

Despite its inability to stimulate TBK1 autophosphorylation as we observed in Fig. 1C, ALTOτ showed the strongest interaction with STING (Fig. 2A) and retained approximately half-maximal interaction with Src (Fig. 2B), indicating it remains structurally and functionally competent in other respects. ALTO-TBK1 interaction peaked with the ALTOΦ variant, declined substantially with ALTOυ, and was reduced to near-background levels at ALTOτ (Fig. 2C). These data suggest that ALTOτ is properly folded and localized but lacks a domain specifically required for ALTO-TBK1 engagement. Taken together, the TBK1 activation and protein-protein interaction assays identify the 30aa region absent in ALTOτ (aa 121-150) as a critical functional segment. We therefore refer to this region as the Lost in Tau (LIT) domain.

### The LIT domain functions as a dominant negative inhibitor of ALTO-induced TBK1 autophosphorylation

To further investigate whether the LIT region is sufficient to confer TBK1-related activity, we tested whether the domain alone could recapitulate or inhibit full-length ALTO function. We first cloned the LIT sequence into the NanoBiT vectors for use in sensitive live cell interaction assays, then subcloned the same reading frame with the addition of the N-terminal LgBiT tag into the pcDNA4c vector, adding an Xpress tag and placing the construct under the control of a highly active CMV promoter for applications requiring more robust expression (Fig. 3A).

**Figure 3:**
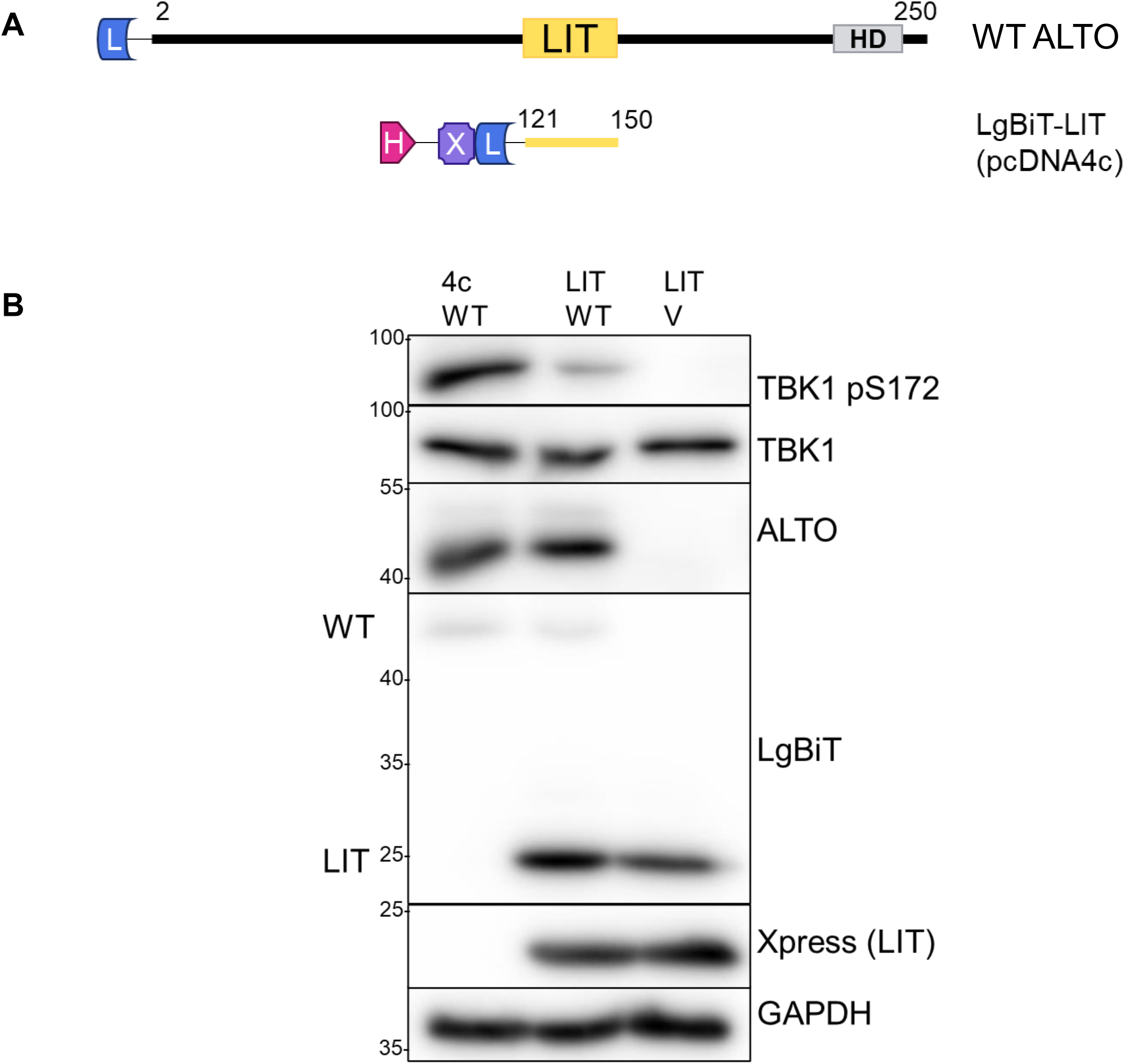
LIT alone is not sufficient to stimulate TBK1 autophosphorylation, but competes with WT ALTO to reduce TBK1 activation. A) Schematic diagram of WT ALTO compared to Xpress-LgBiT-Lost In Tau (LIT) expressed from the pcDNA4c vector. B) Western blots of HEK 293T cells transfected with WT ALTO and LIT or matching vector controls. Primary antibodies are listed on the right; left side labels differentiate multiple bands on the same blot.

NanoBiT protein-protein interaction studies of LIT’s interactions with wild-type (WT) ALTO, STING, Src, and TBK1 were initiated using the standard LgBiT-LIT construct; however, we observed substantial differences in expression levels between WT ALTO and LIT, as well as high variability and background luminescence readings in both experimental and negative control conditions (Fig. S4). Because higher LIT expression would also produce proportionally higher background signal, we concluded that the NanoBiT platform was not suitable for reliably examining LIT interactions.

Since the LIT domain appeared to be critical for ALTO-TBK1 interactions (Figs. 1C, 2C), we hypothesized that it might act as a dominant negative inhibitor by competitively interfering with TBK1 activation by WT protein. To test this, we co-transfected our Xpress-LgBiT-LIT construct with WT ALTO and assessed TBK1 autophosphorylation as well as total TBK1 protein levels. As expected, modest expression of WT ALTO alone strongly stimulated TBK1 autophosphorylation, whereas even much higher expression of Xpress-LgBiT-LIT failed to do so (Fig. 3B). Co-transfection of the two constructs showed a marked decrease in TBK1 autophosphorylation, confirming a dominant negative phenotype.

An alternate ALTO mutant containing the N-terminal 150 aa of ALTO—terminating with the LIT region—was tested as a more stable truncation construct than LIT alone, without requiring a bulky N-terminal tag (Fig. S5). This variant, termed ALTO150, was stably and detectably expressed; however, it had only a modest inhibitory effect on TBK1 autophosphorylation when co-transfected with WT ALTO. Co-transfection also resulted in a modest increase in WT ALTO abundance, suggesting that the N-terminal domain of ALTO mediates its dimerization, which in turn contributes to protein stabilization. While ALTO150 exhibited some dominant negative activity, its stabilization of WT ALTO may offset this effect, creating a self-limiting interaction that blunts its ability to fully suppress WT ALTO function.

### Deletion of the LIT domain abolishes TBK1 activation without disrupting ALTO structure, oligomerization, or other partner binding

While truncation mutants helped define the boundaries of the LIT domain, we next asked whether targeted deletion of this region would phenocopy the loss of function seen in ALTOτ. To determine whether the observed loss of interaction and TBK1 stimulation was due to a sequence-specific function of the LIT region or simply a consequence of reduced length or flexibility, we generated an ALTO variant lacking the LIT coding sequence, termed ALTO deletion of LIT (ALTOdelLIT) (Fig. 4A). The removal of a random 30aa segment was not expected to produce a phenotype, as a deletion of this length in the ALTOΨ truncation did not impair TBK1 autophosphorylation or interaction (Figs. 1C, 2C).

**Figure 4:**
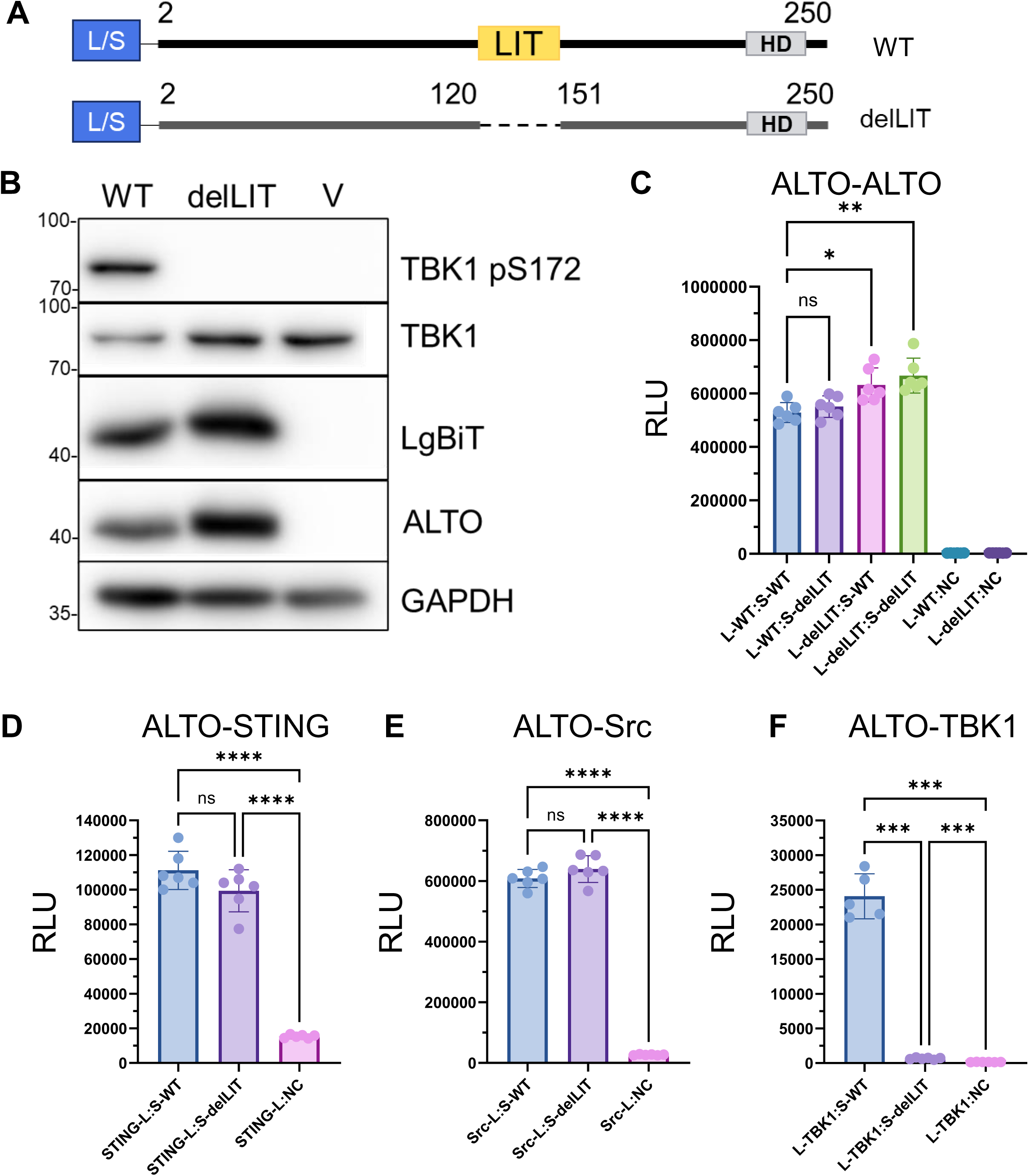
Loss of TBK1 autophosphorylation stimulatory capacity is sequence-specific. (A) Schematic diagram of tagged WT ALTO (top) and ALTO deletion of LIT (ALTOdelLIT, bottom). The location of the LIT region is highlighted by a yellow box. (B) HEK 293T cells were transfected with plasmids carrying either wildtype L-ALTO or L-ALTOdelLIT. Lysates were harvested 24hrs post-transfection and immunoblotted with the indicated antibodies (C-F) HEK 293T were transfected with pairs of plasmids carrying tagged WT ALTO or ALTOdelLIT, as well as tagged STING (D), Src (E), and TBK1 (F). At 20 hours (ALTO-TBK1) or 24 hours (ALTO-STING and ALTO-Src) post-transfection, luminescence was measured. Points indicate biological replicate wells, bars indicate means, and error bars indicate standard deviations. L-protein and protein-S (and similar) indicate the named protein tagged with the LgBiT at its N-terminus or SmBiT at its C-terminus, respectively. All analyses shown are Brown-Forsyth and Welch’s one-way ANOVA, ****p<0.0001; ***p<0.001; **p<0.01; *p<0.05

Consistent with previous results, LgBiT-tagged WT ALTO robustly stimulated TBK1 autophosphorylation and degradation in transfected HEK 293T cells. In contrast, ALTOdelLIT failed to induce either autophosphorylation or degradation of TBK1, despite comparable protein expression levels (Fig. 4B). NanoBiT interaction assays confirmed that this loss of function was not due to impaired ALTO oligomerization, as ALTOdelLIT retained equal or even enhanced capability to oligomerize with itself and with WT ALTO (Fig. 4C). Similarly, interactions between ALTOdelLIT and STING (Fig. 4D), as well as ALTOdelLIT and Src (Fig.4E), were not significantly different from those of WT ALTO. Interaction between ALTOdelLIT and TBK1, however, was reduced by more than 30-fold compared to WT ALTO (Fig. 4F), indicating a specific requirement for the LIT region in TBK1 engagement. Protein expression across all interaction assays was largely equal, further supporting the reliability of these findings (Fig. S6).

The preserved interactions between ALTOdelLIT and ALTO, STING, and Src suggest that ALTOdelLIT is structurally and functionally intact, and that its loss of TBK1 engagement reflects a specific requirement for the LIT region.

The LIT region lies within ALTO’s N-terminal intrinsically disordered region (IDR), and as such, no confident structure-based predictions can be made about its function or (Fig. 1A). To further investigate whether specific functional domain(s) of ALTO within the LIT region might drive the ALTO-TBK1 interaction, we analyzed its amino acid sequence for short linear motifs (SLiMs), also known as eukaryotic linear motifs (ELMs), using the ELM Resource prediction tool (http://elm.eu.org/search.html). The analysis was performed without specifying a subcellular compartment, as ALTO’s subcellular localization remains poorly defined. The resulting predictions were then filtered to include only those located within or adjacent to the LIT region (Fig. S7). Although SLiMs are inherently degenerate and therefore not considered high-confidence predictions when found, no motifs corresponding to known TBK1-binding sites or interaction domains with other signal complex components were identified.

### The LIT domain does not influence ALTO subcellular localization or trafficking

Given that ALTOdelLIT retains structural integrity and continues to interact with Src, STING, and itself, we next hypothesized that differences in intracellular trafficking might account for the phenotypic divergence between WT ALTO and ALTOdelLIT. To test this, we generated human dermal fibroblasts (HDFs) expressing either WT ALTO or ALTOdelLIT under a doxycycline (Dox) inducible promoter (inALTOWT and inALTOdelLIT HDFs, respectively). After several hours of Dox induction, cells were fixed and immunostained for ALTO and various intracellular membrane markers. Substantial colocalization was observed only with the cis-Golgi marker GM130, where both WT ALTO and ALTOdelLIT appeared to accumulate during early expression (Fig. 5A). At later time points, both proteins were observed to traffic outward from the Golgi, potentially via the secretory pathway (Fig. S8). Further immunofluorescence analysis revealed partial colocalization of distal ALTO and ALTOdelLIT foci with the early endosome marker Rab5 (Fig. 5B), suggesting ALTO may transiently localize to the plasma membrane and re-enter the cell via the endocytic pathway, a finding consistent with prior reports of ALTO co-fractionating with lipid raft markers(31).

**Figure 5:**
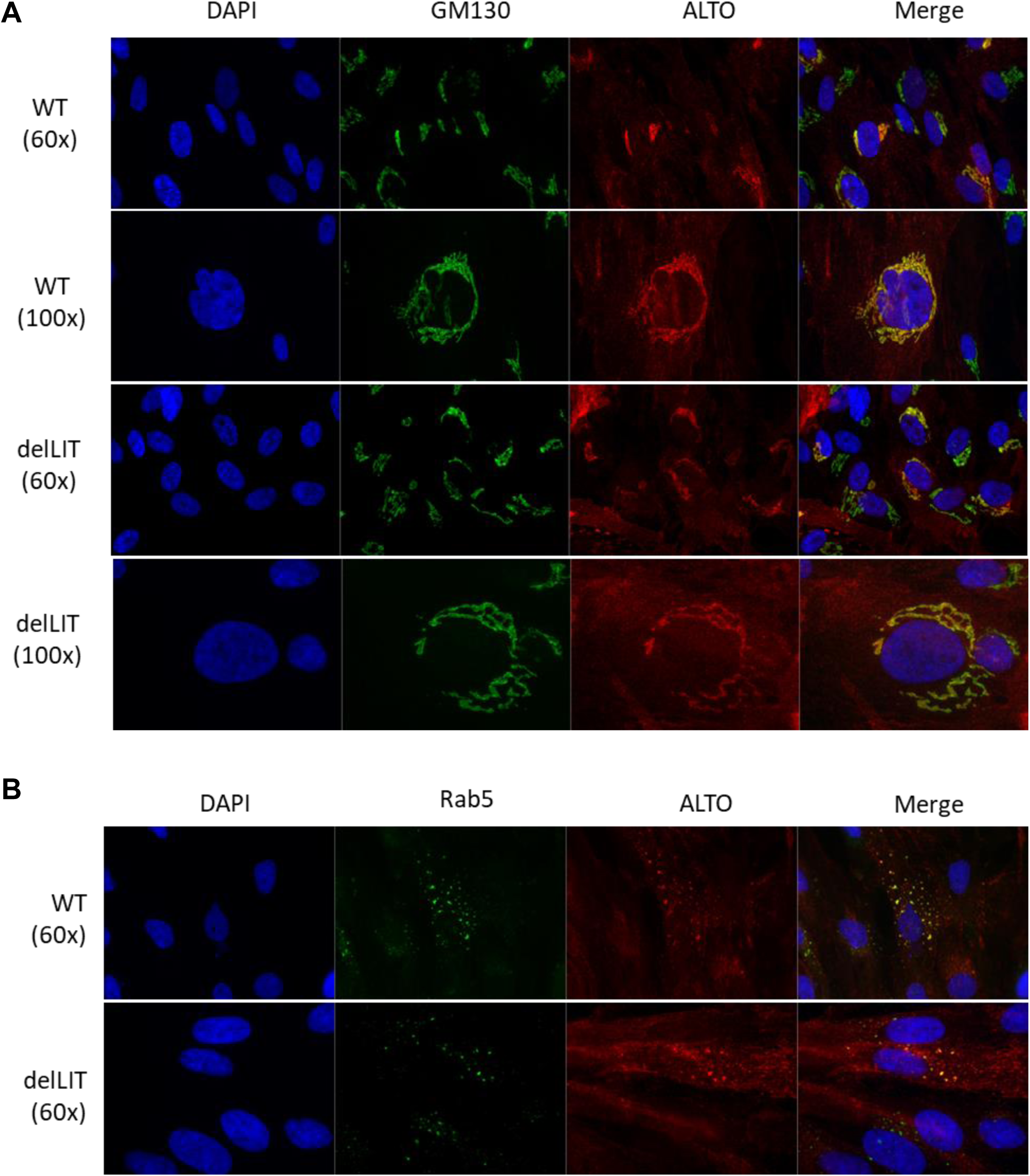
WT ALTO and ALTOdelLIT traffic through the Golgi. (A) inALTO and inALTOdelLIT HDFs induced for 4 hours prior to fixation and immunofluorescent stained for ALTO and the cis-Golgi marker GM130. (B) Representative immunofluorescence images of inALTO and inALTOdelLIT HDFs fixed at 16 hours post-induction and stained for ALTO and the early endosome marker Rab5.

Although we did not identify a specific motif or altered trafficking pattern that could explain the loss of ALTO-TBK1 interaction in the absence of the LIT domain, these findings suggest that ALTO’s trafficking may involve dynamic cycling through the secretory and endocytic pathways. Further investigation into ALTO’s trafficking and intracellular lifecycle may provide important insights. Despite the absence of clear structural or localization changes, our data demonstrate that the LIT domain is specifically required for ALTO-induced TBK1 autophosphorylation.

### The LIT domain is essential for ALTO-mediated suppression of MCPyV replication in infected cells

Finally, to place our protein-level observations into their full viral context, we evaluated whether WT ALTO and ALTOdelLIT could suppress MCPyV replication by using our HDF model of early MCPyV infection(40). Donor-matched inALTOWT and inALTOdelLIT HDFs (Fig. 6A) were infected with equal titers of ALTO_null_ MCPyV, a genetically intact virus containing a point mutation in the ALTO start codon that eliminates ALTO expression(29). Following infection, cells were either mock-induced or induced to express their respective ALTO variant, and viral replication was assessed five days post-infection.

**Figure 6:**
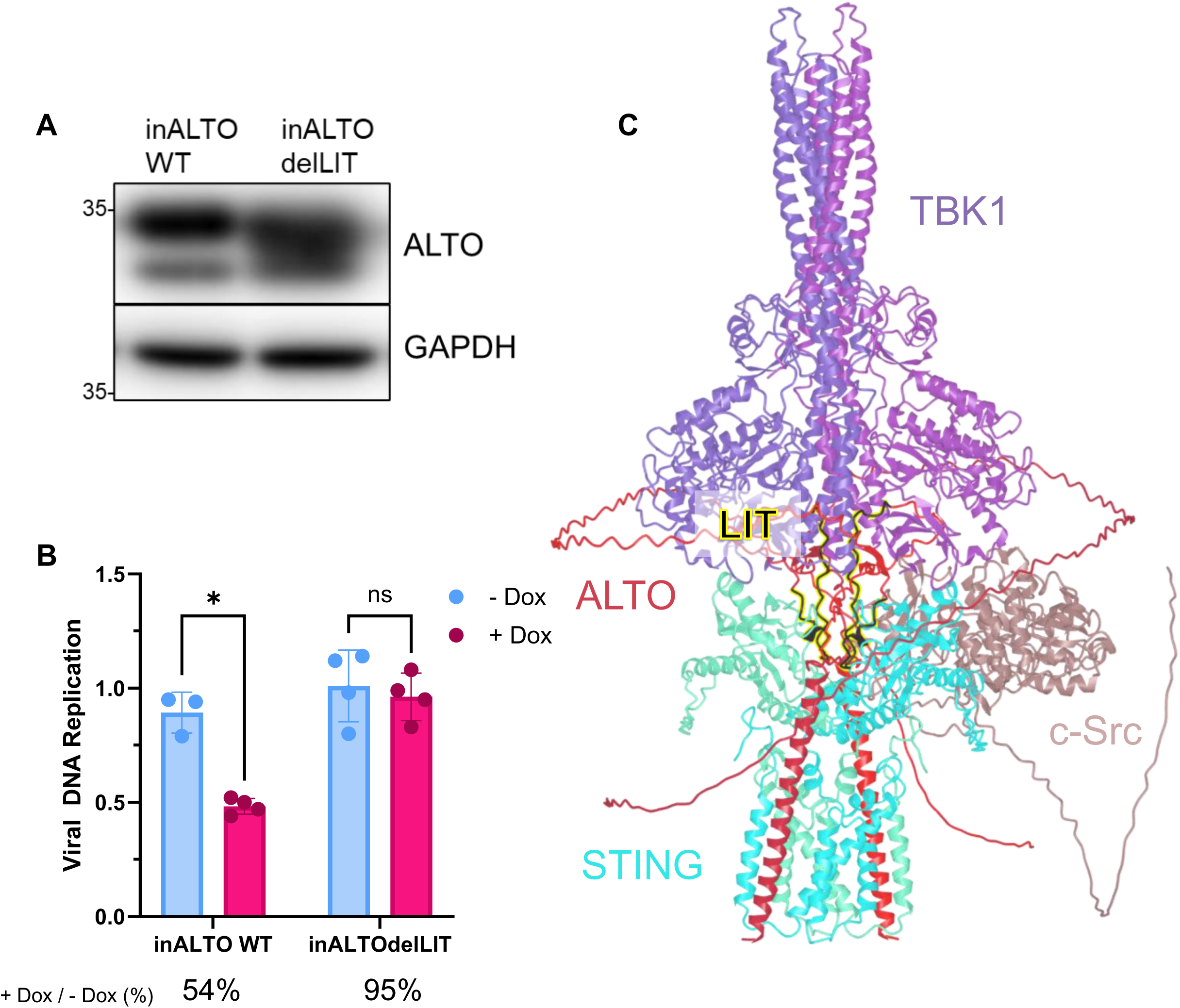
The LIT domain is essential for ALTO-mediated suppression of MCPyV replication in infected cells. (A) Western blots of HDFs inducibly expressing ALTOdelLIT or WT ALTO. Lysates were collected 16 h post-induction. (B) qPCR analysis of viral DNA extracted 5 days post-infection from HDFs inducibly expressing WT ALTO or ALTOdelLIT, following infection with equal titers of ALTO_null_ MCPyV. Analysis is normalized to cellular GAPDH for each cell type. Points represent biological replicate wells averaged from three technical replicates each. Analyses are multiple unpaired t-tests with Welch’s correction, *p<0.05. (C) Predicted tertiary structure modeling of the ALTO/STING/TBK1/Src complex, with the LIT domain highlighted at center. Full-length canonical amino acid sequences for each protein were used with AlphaFold3 server (update 2025.06.10) with no post-translational modifications, default parameters, random seeding.

As we previously reported, ALTO_null_ virus replicates significantly more efficiently in uninduced HDFs compared to HDFs induced to express WT ALTO. In contrast, induction of ALTOdelLIT failed to significantly reduce viral replication (Fig. 6B). Because the suppression of viral replication by ALTO is known to occur through a TBK1-driven negative feedback mechanism(29), this loss of function demonstrates that the LIT domain is required for TBK1 stimulation by ALTO during viral infection.

## Discussion

MCPyV remains the only conclusively identified oncogenic human polyomavirus. Despite its widespread presence in the general population and its clear role in the pathogenesis of MCC, major gaps persist in our understanding of key aspects of MCPyV biology. Among the most critical of these are the mechanisms by which MCPyV establishes persistent infection, maintains latency, and ultimately exits latency to drive oncogenesis.

This study builds on our previous work proposing an MCPyV latency mechanism involving ALTO-mediated modulation of the STING–TBK1 axis. To further define this mechanism, we used a progressive N-terminal truncation approach spanning ALTO’s intrinsically disordered region (IDR) and alpha-helical domain to identify regions critical for TBK1 activation (Figs. 1A, 1B). Through this strategy, we discovered a short, previously uncharacterized region of ALTO that is specifically required for TBK1 stimulation. The ALTOτ mutant, which retains only the C-terminal 100 amino acids of ALTO (aa 151–250), was the first truncation to completely lose the ability to stimulate TBK1 autophosphorylation in cellulo (Fig. 1C) and showed almost no detectable interaction with TBK1 in protein–protein interaction assays (Fig. 2C). In contrast, its immediate upstream predecessor, ALTOυ, retained both activity and binding, indicating that the 30 amino acids present in ALTOυ but absent in ALTOτ (aa 121–150) are essential for this function. We therefore designated this region the LIT (Lost in Tau) domain. Located within ALTO’s IDR, the LIT domain is predicted to localize at the interface of ALTO, STING, TBK1, and Src within the multi-protein signal complex (Fig. 6C).

When expressed in isolation, the LIT domain acted as a dominant negative inhibitor of WT ALTO, competitively blocking TBK1 activation (Fig. 3B). To confirm that the LIT domain is necessary for ALTO’s immune modulatory activity, we engineered a deletion construct (ALTOdelLIT) and found that it failed to stimulate TBK1 autophosphorylation (Fig. 4B). This loss of function appears to be sequence-specific: ALTOdelLIT maintained expression, proper folding, and interactions with other signal complex proteins such as STING and Src, but showed a dramatic reduction in TBK1 binding (Fig. 4F). This loss was not attributable to the absence of a canonical TBK1-interaction motif (Fig. S7) or to altered protein trafficking, as ALTO and ALTOdelLIT both localized to the cis-Golgi at early time points and trafficked through both the secretory and endocytic pathways (Figs. 5A, 5B).

Finally, we demonstrated that the LIT domain is functionally required for TBK1-mediated suppression of MCPyV replication during early infection (Fig. 6A). Notably, ALTOdelLIT induction failed to reduce viral replication to the extent observed with direct pharmacological inhibition of TBK1(29) in our prior study. These results support the conclusion that ALTO, through the LIT domain, plays a central role in modulating host–virus interactions during both early and persistent infection.

The identification of a novel immune interaction domain within ALTO’s IDR builds on prior reports implicating other regions of ALTO in Src-family kinase binding(29, 31), PLCγ1 signaling, and NFκB modulation(31, 32). Collectively, these studies revise the earlier view of ALTO as a dispensable accessory protein and instead establish it as a key viral effector that actively shapes innate immune responses through structurally dynamic interactions. As ALTO-encoding polyomaviruses continue to emerge as a phylogenetic subset of mammalian polyomaviruses(32), it will be important to assess whether the LIT domain is conserved across species or represents a specialized adaptation of MCPyV to its human host. Given that many polyomaviruses establish persistent infections, conservation of LIT-like motifs would support the idea that latency is a regulated process driven by active host–virus interactions, rather than a passive consequence of viral quiescence.

MCPyV ALTO is not unique among viral or oncogenic proteins in its use of intrinsic disorder to regulate complex signaling networks. Other viral effectors, including HPV E7(41) and KSHV ORF57(42), also use disordered regions to engage host pathways with high flexibility and specificity. As the functional importance of intrinsically disordered viral proteins becomes more widely appreciated, new opportunities are emerging to map these regions and understand their mechanistic roles in immune evasion, persistence, and oncogenesis(43–45).

Several key questions remain regarding the lifecycle and regulation of ALTO during MCPyV infection. Although we observed initial accumulation of ALTO at the cis-Golgi, followed by trafficking through secretory and endocytic pathways (Figs. 5, S8), and prior studies have reported association with lipid rafts(31), the complete subcellular dynamics of ALTO, particularly during long-term infection, are not yet well defined. Given the known importance of spatial regulation for many signaling proteins, a deeper understanding of ALTO’s localization and trafficking may reveal new insights into its diverse functions and regulatory partners.

A particularly intriguing and unresolved observation is the frequent loss of ALTO expression in virus-positive MCC, even in cases where the ORF is retained(32). While some loss may result from stochastic truncation events during viral integration, ALTO protein is undetectable in several virus-positive MCC lines that retain its intact coding sequence. From an evolutionary perspective, this may be a form of immune evasion. ALTO is highly immunostimulatory, and its continued expression after integration could trigger immune clearance of the host cell. Thus, the absence of ALTO in MCC tumors may reflect a form of survivorship bias. Only those cells that have lost ALTO expression may escape immune detection and progress to cancer. Since ALTO can be robustly expressed even in the absence of all other viral proteins, this suggests that host-mediated regulatory mechanisms, rather than viral factors, may suppress ALTO post-integration. Identifying such factors could reveal novel therapeutic targets or biomarkers for MCC progression.

Finally, the discovery of a compact domain within ALTO that selectively activates TBK1 opens the door to potential translational applications. ALTO itself or a small molecule mimic of its TBK1 activating domain could serve as a noncanonical TBK1 agonist for boosting antiviral immunity in immunocompromised individuals, potentially suppressing MCPyV replication and reducing cancer risk. Beyond direct antiviral applications, TBK1 is a known druggable target for a variety of conditions associated with its dysregulation, as well as a target of interest in the development of novel cancer immunotherapies(46–48). Identification and characterization of the noncanonical mechanism by which ALTO is able to stimulate TBK1 autophosphorylation could also enable the development of novel agonists and antagonists. Conversely, the LIT domain itself, or derivatives thereof, could act as dominant negative inhibitor of WT ALTO by blocking its ability to activate TBK1. This raises the possibility of repurposing LIT-mimicking compounds to modulate ALTO-TBK1 signaling in MCPyV-associated conditions, such as latency control or early-stage infection.

## Materials and methods

### Cell culture

All cells were maintained at 37°C with 5% CO_2_ in a humidified incubator. HEK 293T cells were cultured in Dulbecco’s Modified Eagle Medium (DMEM, Gibco) supplemented with 10% fetal bovine serum (FBS, Cytiva). HDFs were isolated from neonatal foreskin as previously described(49) and maintained in DMEM supplemented with 10% tetracycline-screened FBS (Cytiva).

### Chemicals and reagents

Doxycycline (Dox) and puromycin were dissolved in sterile water to stock concentrations of 0.5 mg/mL and 10 mg/mL, respectively, and stored at −20°C. Unless otherwise noted in the text or figure legends, Dox was used at a final concentration of 0.5 μg/mL. Puromycin was used during selection for stable cell lines as described below. Hexadimethrine bromide (Polybrene, Aldrich) was dissolved in water to a 12 mg/mL stock, sterilized by 0.2 μm syringe filtration, and used fresh for each experiment.

### Recombinant plasmid construction

NanoBiT constructs expressing WT ALTO, TBK1 and STING were generated in our previous study(29). The human c-Src coding sequence (hereafter “Src”) was amplified from the pDEST40-ZXHA-WtSrc plasmid and cloned using XhoI and NheI sites. LgBiT- and SmBiT-tagged truncated ALTO mutants (Ψ, χ, Φ, υ, τ, σ, ρ, ΔTM, and LIT) were PCR-amplified from the pBiT1.1-N_ALTO (L-ALTO) backbone and subcloned into pBiT1.1-N[TK/LgBiT] or pBiT2.1-N[TK/SmBiT] vectors using XhoI and NheI.

Deletion constructs pBiT1.1-N_ALTOdelLIT (L-ALTOdelLIT) and pBiT2.1-N_ALTOdelLIT (S-ALTOdelLIT) were constructed by linear PCR omitting the LIT region, followed by Bpu10I digestion and self-ligation.

Constructs for pcDNA4c-ALTO150 and pcDNA4c-LgBiT-LIT were prepared by amplifying their respective coding sequences from their respective NanoBiT plasmids and subcloning into the pcDNA4c vector (Invitrogen) using BamHI and XhoI or EcoRI, respectively.

The pTRIPz-RFP vector (Open Biosystems) and pTRIPz-ALTO construct were generated as previously described(29). pTRIPz-ALTOdelLIT was derived from pBiT1.1-N_ALTOdelLIT and cloned into the pTRIPz backbone using the AgeI and MluI sites to replace the RFP cassette. All primer sequences are provided in Supplemental Table 1.

### Structural prediction

Structural modeling of WT ALTO (Fig. 1A) was performed as previously described(29, 35). Progressive ALTO truncation mutants and the STING-TBK1-ALTO-Src complex were modeled using the AlphaFold3 server (builds 2025.5.23 and 2025.06.10) with default settings(34). Models were visualized and recolored using NIH iCn3D(50, 51).

### Generation of inducible stable HDFs

Inducible HDF lines were generated as previously described(27, 49). Briefly, lentiviral particles were produced in HEK 293T cells by co-transfecting pTRIPz transfer plasmids (encoding WT ALTO or ALTOdelLIT), psPAX, and pMD2.G using Lipofectamine 2000 (Invitrogen). Supernatants were collected 48 h post-transfection, centrifuged, 0.2 μm filtered, and supplemented with 6 μg/mL Polybrene prior to transduction of early-passage HDFs (≤2 passages post-isolation). After 48 h, media was replaced, and cells were selected with 4 µg/mL puromycin for 5 days, followed by maintenance in 1 µg/mL puromycin.

### NanoBiT protein–protein interaction assays

HEK 293T cells were seeded into opaque white 96-well plates (3×10⁴ cells/well) in 100 μL DMEM + 10% FBS. Warm PBS was added to outer wells and between wells to reduce edge effects. The following day, cells were transfected with 100 ng total plasmid DNA per well (50 ng each of LgBiT and SmBiT constructs or SmBiT control) using FuGENE HD (Promega) in Opti-MEM I (Gibco). After 20–24 h incubation (20 h for TBK1 assays), media was replaced with fresh Opti-MEM I ∼30 min before reading. NanoGlo Live Cell Substrate (Promega) was diluted to 5X in Live Cell Buffer and added under low-light conditions. Luminescence was measured using a Luminoskan Ascent (Thermo) every 15 minutes for 90 minutes. Representative timepoints were selected based on signal strength and stability.

### Western blotting

Cells were lysed in ice-cold buffer (10 mM HEPES pH 7.5, 500 mM NaCl, 1 mM EDTA, 1 mM DTT, 0.5% Triton X-100) supplemented with PhosSTOP (Roche) and cOmplete protease inhibitors (Roche). Lysis proceeded on ice for 30–60 min with intermittent vortexing. Lysates were cleared by centrifugation at 12,000 × g for 10 min at 4°C. Protein concentration was determined by Bradford assay. Samples were mixed with Laemmli buffer, boiled for ≥2 min, flash frozen, and stored at −80°C. Proteins were resolved by SDS-PAGE and transferred to 0.45 μm PVDF membranes. Blots were blocked and incubated overnight at 4°C with primary antibodies diluted in 5% milk in PBST (0.1% Tween-20), followed by secondary antibodies for 1 h at room temperature. Signal was developed using SuperSignal West Pico PLUS (Thermo) and imaged on an Amersham Imager 600 or 680 (GE Healthcare). Primary antibodies: anti-GAPDH (1:5,000, CST 2118S), anti-TBK1 (1:1,000, CST 3504S), anti-phospho-TBK1 S172 (1:1,000, CST 5483S), anti-ALTO (1:10,000, courtesy of the Galloway Lab), anti-STING (1:1,000, CST 13647S), anti-LgBiT (1:500, Promega N710A), anti-Xpress (1:1,000, Invitrogen 44-0528), anti-Src (1:1,000, CST 2109S). Secondary antibodies: HRP-conjugated anti-rabbit IgG (1:2,500, CST 7074S), anti-mouse IgG (1:2,500, CST 7076S).

### Immunofluorescent staining

HDF-inALTO and -inALTOdelLIT cells were seeded on coverslips and treated with Dox (0.5 μg/mL) for durations indicated in the figures. Cells were fixed in 3% paraformaldehyde (PBS, 20 min), permeabilized in 0.5% Triton X-100 / 3% BSA (1 h, RT), and incubated with primary antibodies in the same buffer for 1 h in humidified chambers. Following PBS washes, secondary antibodies with DAPI were applied under low-light conditions. Cells were imaged using an Olympus IX81 inverted microscope. Primary antibodies: anti-ALTO (1:1,000, courtesy of the Galloway Lab), anti-Rab5 (1:200, CST 3547T), anti-GM130 (1:200, BD 610822). Secondary antibodies: AlexaFluor 594-conjugated goat anti-rabbit IgG (1:500, Invitrogen A11012), AlexaFluor 488-conjugated goat anti-mouse IgG (1:500, Invitrogen A11029).

### MCPyV infection

Infections using ALTO_null_ MCPyV were performed in HDF-inALTO and -inALTOdelLIT lines as previously described(29). Viral replication was assessed by qPCR following published methods(29).

### Statistical analysis

All statistical analyses were performed using GraphPad Prism. For single comparisons, unpaired two-tailed t-tests with Welch’s correction were used. For multiple group comparisons, Brown–Forsythe and Welch’s ANOVA tests were applied where indicated. Significance thresholds were: P < 0.05 (*), P < 0.01 (**), P < 0.001 (***). Confidence intervals were set at 95%.

---

**Supplemental Table 1:**
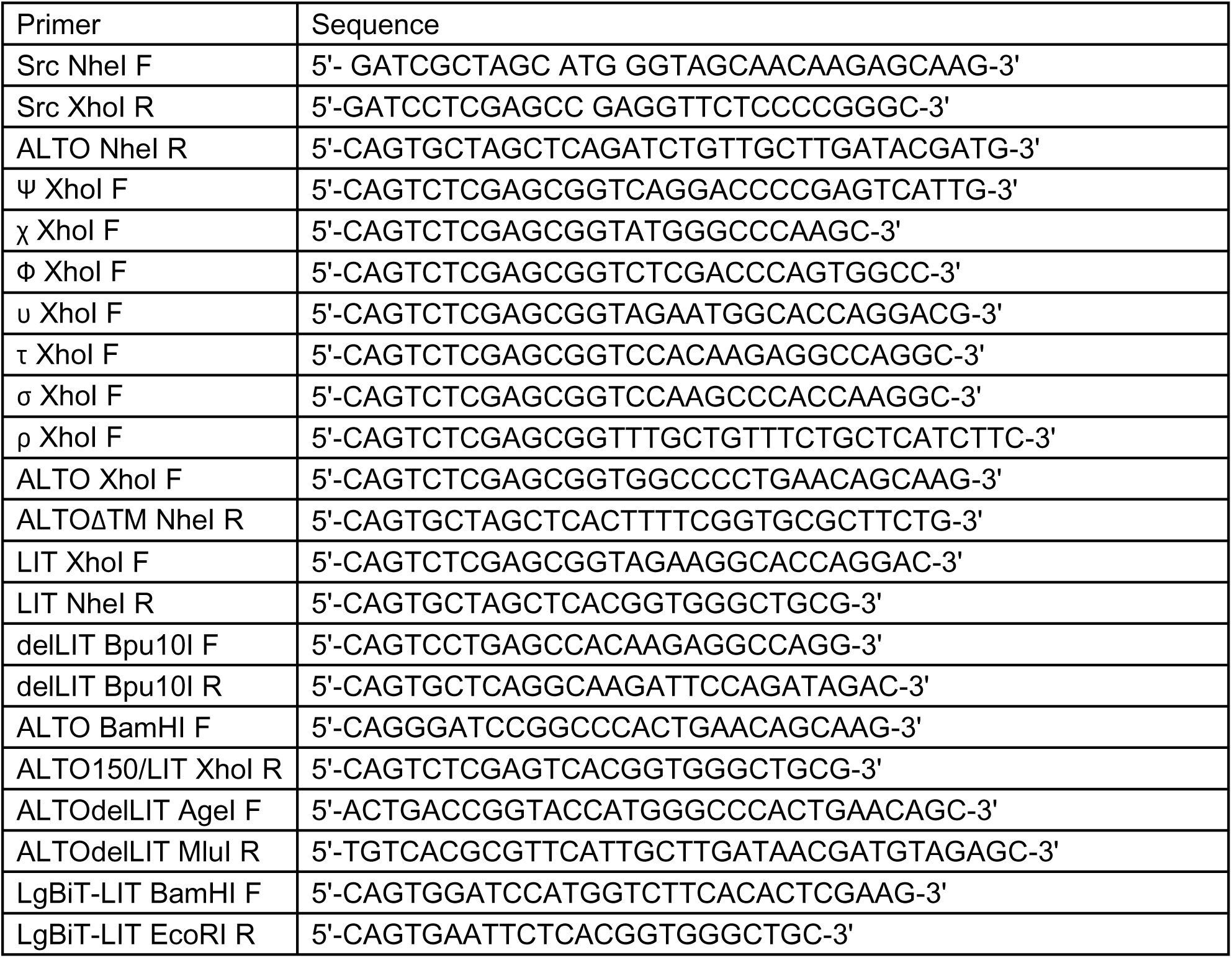

## Acknowledgments

We thank Dr. Denise Galloway for generously providing the anti-ALTO antibody. We are also grateful to the members of our laboratories for their insightful discussions and ongoing support. This work was supported by the National Institutes of Health through Grants R01CA187718, R21CA267803, and R01CA284690 awarded to J.Y., as well as by the National Cancer Institute through P01CA281867 Project 3 to J.Y.. The funders had no role in study design, data collection, interpretation, or submission for publication.

---

**Figure S1:**
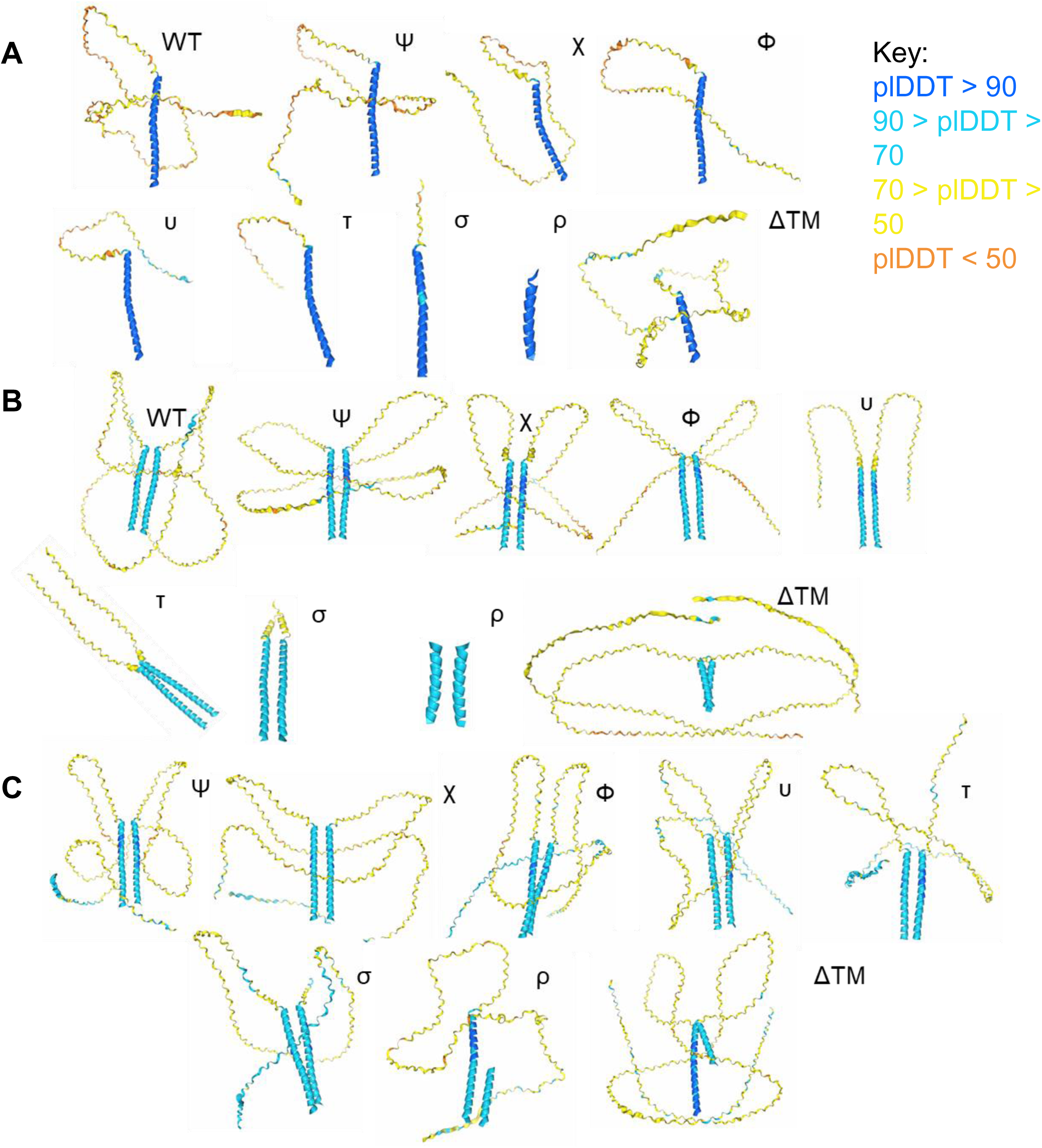
Predictive tertiary structures of progressive ALTO truncation mutants. Amino acid sequences of progressive ALTO truncation mutants were used with the 2025.5.23 Update of the AlphaFold3 server platform (default settings, random seeding). All structures have a low-confidence intrinsically disordered N-terminal region, and a relatively higher-confidence C-terminal alpha helical structure, which is maintained through all truncations. All progressive ALTO truncations are shown in (A) monomeric, (B) homodimeric, and (C) WT ALTO-paired heterodimeric forms. Colors indicate AlphaFold calculated predicted local distance difference test (plDDT) confidence scores according to the supplied key in (A).

**Figure S2:**
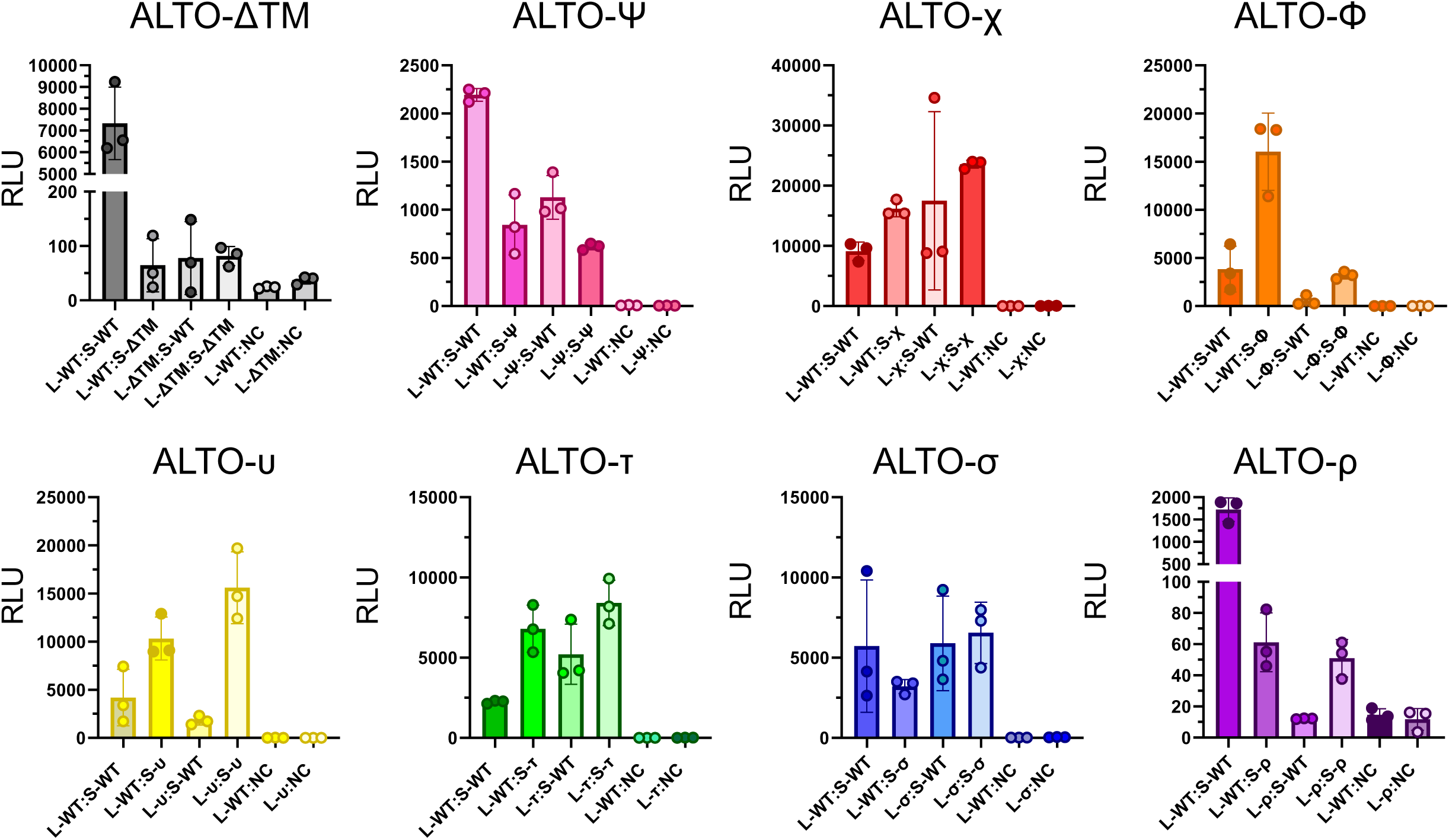
Progressive ALTO truncation mutants retain homo-and hetero-oligomerization potential. Representative NanoBiT luminescence readings of HEK 293T cells transfected with pairs of plasmids carrying NanoBiT-tagged WT ALTO or a truncation mutant, or the Small BiT negative control construct. Luminescence was measured at 24hrs post-transfection and reported in Relative Luminescence Units (RLU). Points indicate biological replicate wells, bars indicate means, and error bars indicate standard deviations. L-protein and protein-S (and similar) indicate the named protein tagged with the LgBiT at its N-terminus or SmBiT at its C-terminus, respectively.

**Figure S3:**
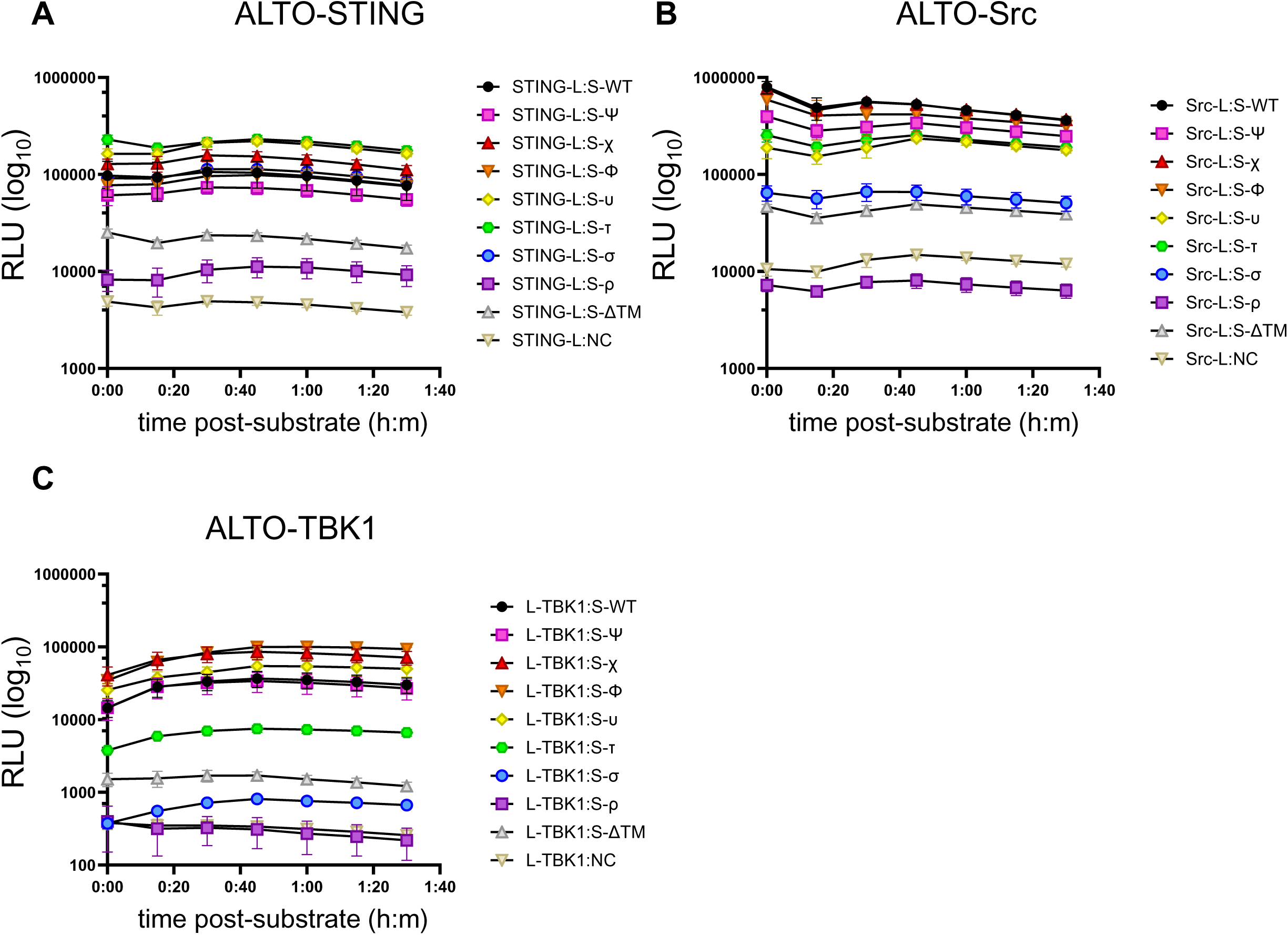
Kinetic curves of progressive ALTO truncation mutant interactions show relative interaction stability over time. Kinetic luminescence readings of HEK 293T cells 20hrs (ALTO-TBK1) or 24hrs (ALTO-STING and ALTO-Src) post-transfection with the indicated NanoBiT interaction pairs. Cells were monitored for 90 minutes after the addition of substrate, with readings recorded every 15min to determine the intensity and stability of protein-protein interactions. Individual points reflect average values of biological replicate wells; error bars indicate standard deviation across replicates at each time point; and trendlines indicate change between averages across timepoints. Single representative timepoints are highlighted in corresponding figures in Fig. 2.

**Figure S4:**
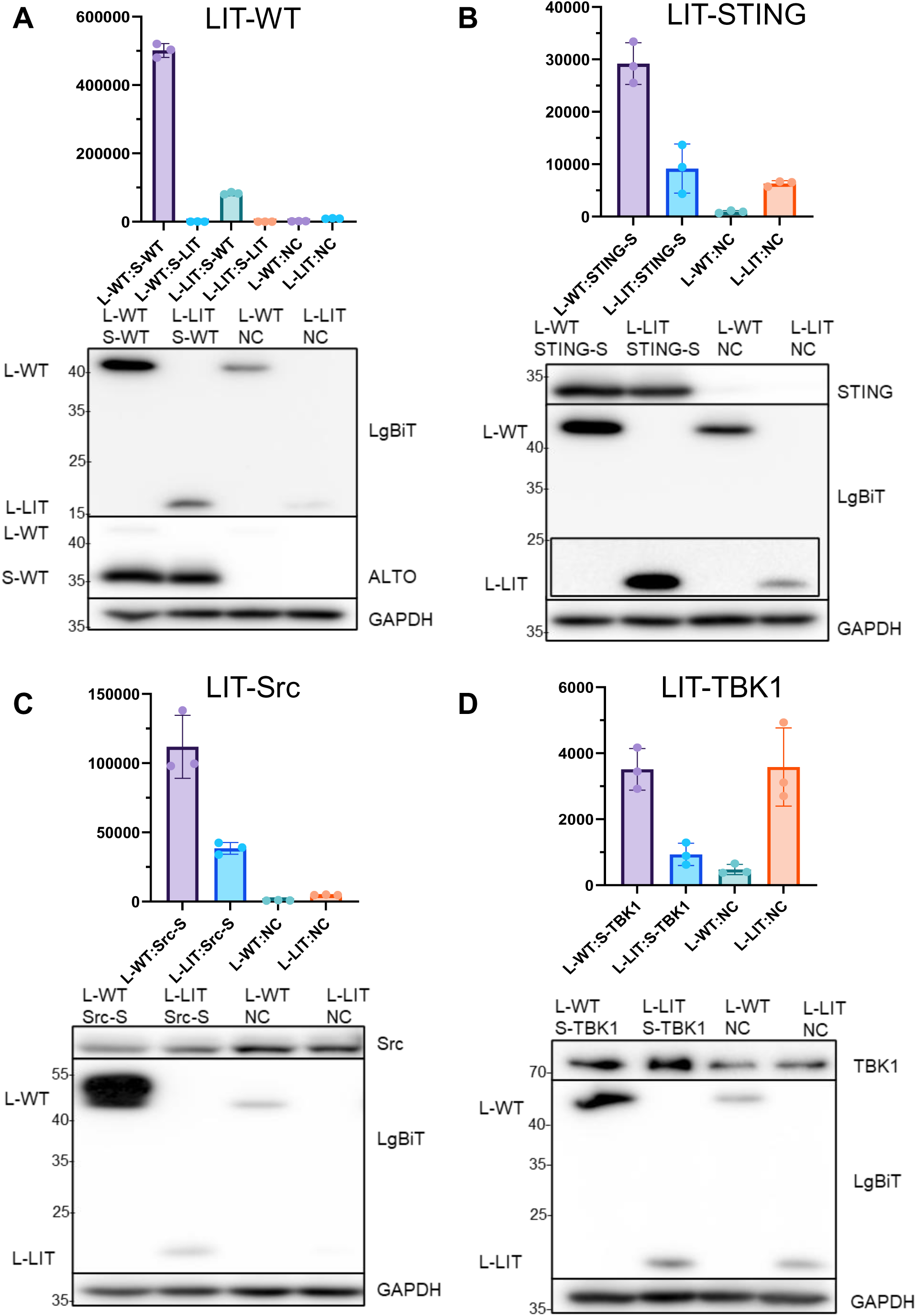
Protein expression in LgBiT-LIT NanoBiT experiments is not sufficiently equal to WT to draw reliable interaction conclusions. Western blots and representative NanoBiT readings of HEK 293T cells co-transfected with tagged LIT and (A) WT ALTO, (B) STING, (C) Src, and (D) TBK1 at 20 hours (TBK1) or 24 hours (all others) post-transfection. Overexposed L-WT bands in (B) were covered to image the L-LIT bands (inset). Primary antibodies are indicated on the right; left-side labels differentiate bands visualized on the same blot.

**Figure S5:**
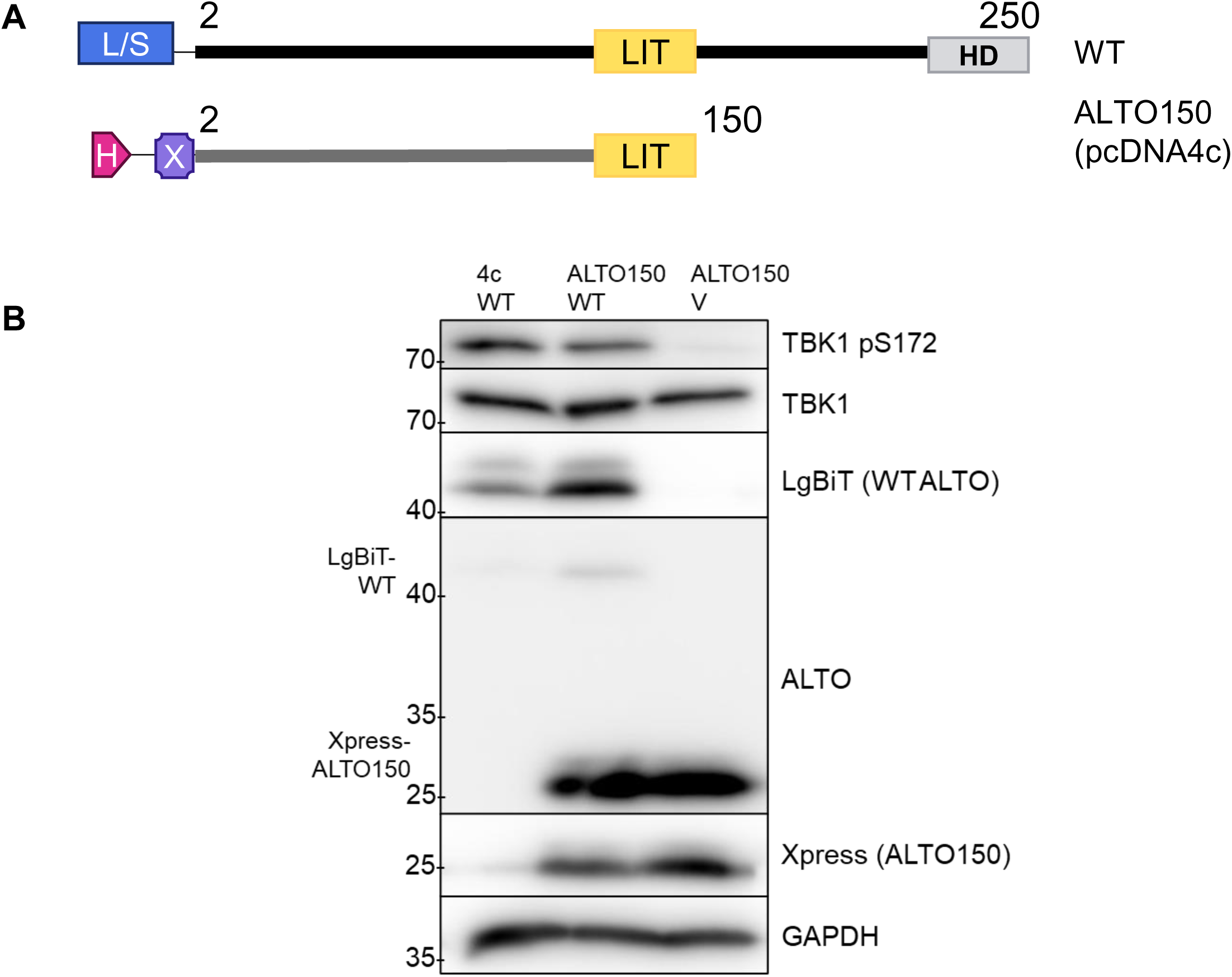
ALTO150 is not sufficient to stimulate TBK1 autophosphorylation but is a partial dominant-negative. (A) Schematic diagram of LgBiT-tagged WT ALTO and 6* His-Xpress-tagged ALTO150 as expressed from the pcDNA4c vector (B) Western blots of lysates from HEK 293T cells 24 hours post-transfection with WT ALTO, ALTO150, or both. Right side labels indicate primary antibodies; left side labels distinguish multiple bands on the same blot.

**Figure S6:**
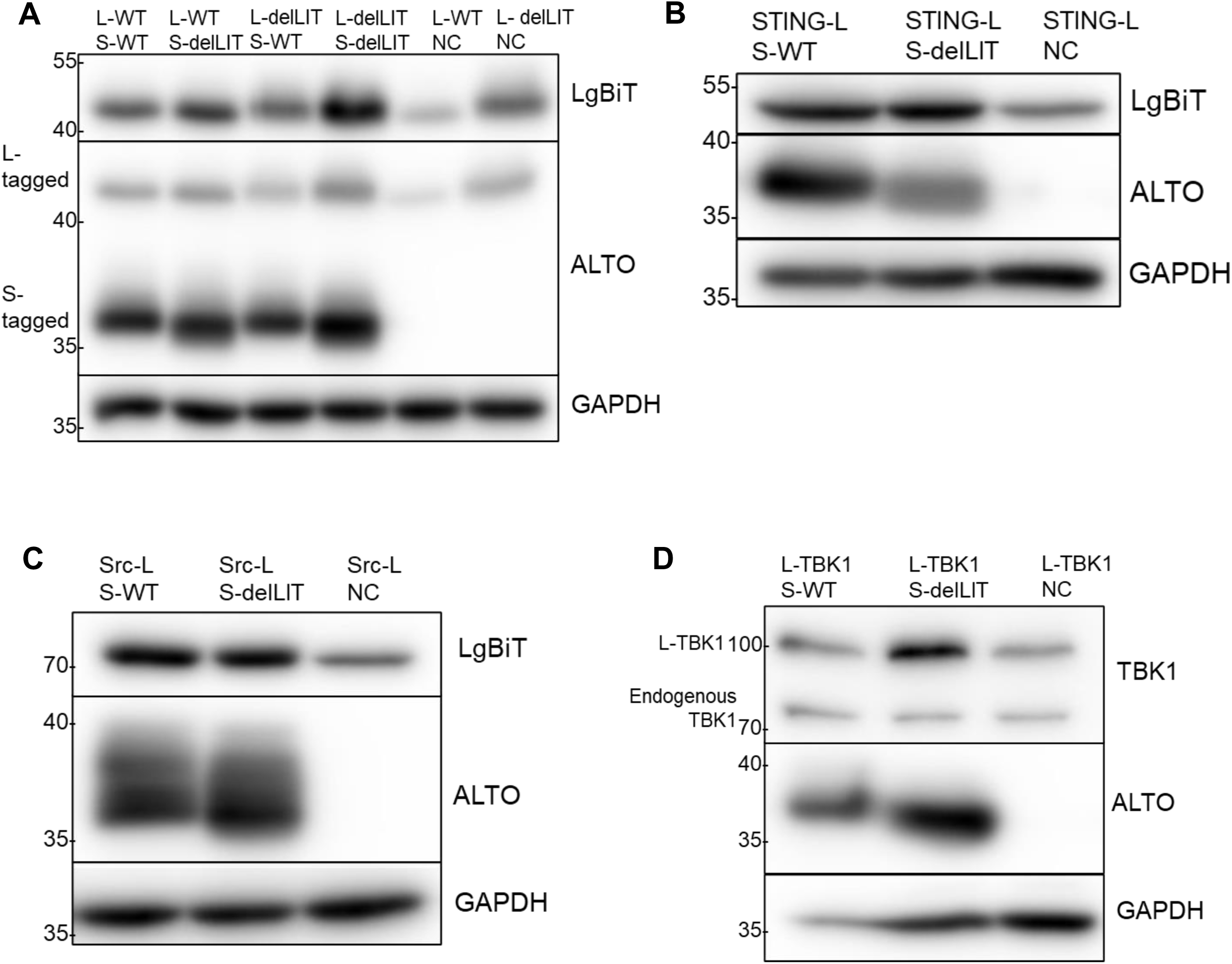
Relatively equal protein expression across variants in ALTOdelLIT NanoBiT experiments supports functional conclusions. Western blots of lysates from HEK 293T cells transfected with indicated NanoBiT pairs, corresponding to Fig. 4. Lysates were harvested at 20hrs (ALTO-TBK1) or 24hrs (all others) post-transfection to assess relative protein expression at the time of luminescence readings. Primary antibodies are listed on the right; labels on the left differentiate multiple bands on the same blot.

**Figure S7:**
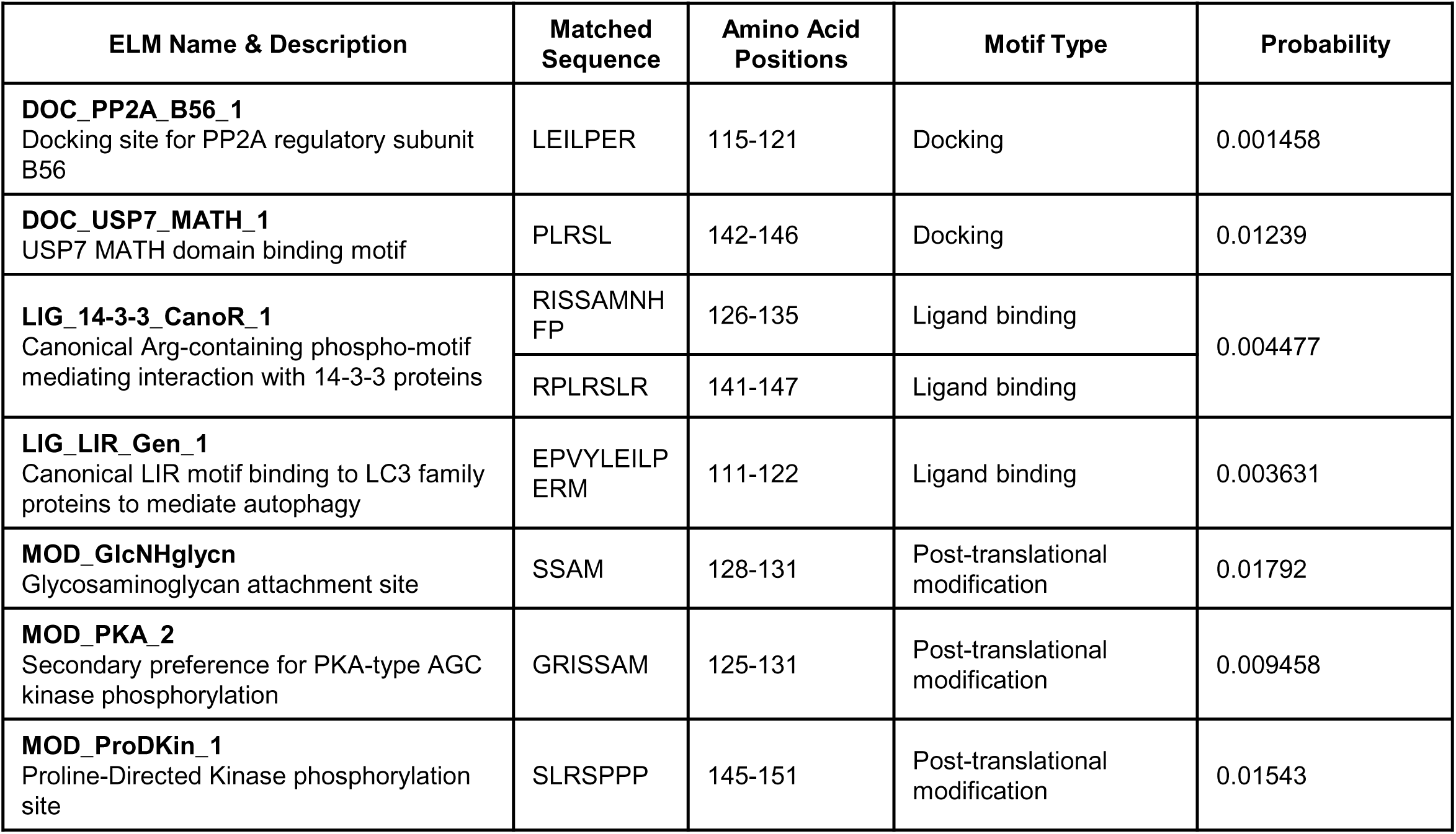
SLiMs do not explain the loss of function in ALTOdelLIT. Full-length amino acid sequence of ALTO was submitted to the ELM database (elm.eu.org) functional search prediction (cell compartment not specified, default parameters, Homo sapiens species context). Results have been filtered to display only motifs which are present within or immediately adjacent to the LIT domain. Probabilities provided by the ELM database reflect the likelihood of the motif to appear by random in any given protein sequence.

**Figure S8:**
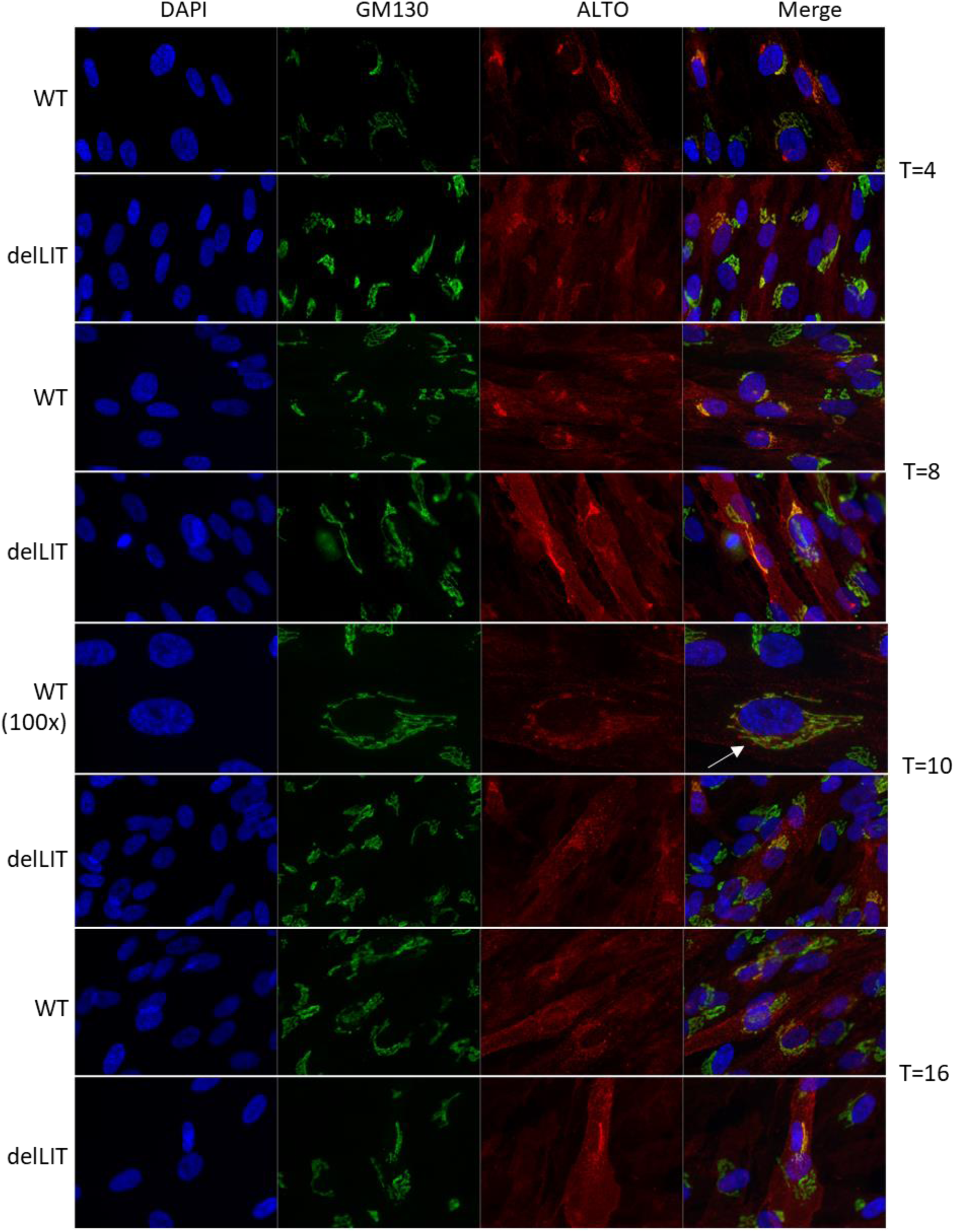
Longitudinal trafficking of WT ALTO and ALTOdelLIT. Representative immunofluorescence images of inALTO and inALTOdelLIT HDFs fixed at the indicated timepoints post-induction. All images taken with the 60x objective unless otherwise indicated. The WT cell shown at 100x at T=10 (white arrow) was particularly noteworthy for the apparent visualization of ALTO punctae leaving the Golgi.

## Notes

### Competing Interest Statement

The authors have declared no competing interest.

### Summary of Updates

Supplementary figures S1, S2, S4, and S5 have been revised to correct for image errors caused during initial pdf conversion.

